# ABEL-FRET bridges the timescale gap in single-molecule measurements of the structural dynamics in the A_2A_ adenosine receptor

**DOI:** 10.1101/2025.07.22.666123

**Authors:** Ivan Maslov, Valentin Borshchevskiy, Vadim Cherezov, Johan Hofkens, Jelle Hendrix, Michael Börsch, Thomas Gensch

## Abstract

The functional complexity of G protein-coupled receptors (GPCRs) arises from their structural dynamics, which span timescales from nanoseconds to minutes. Single-molecule Förster Resonance Energy Transfer (smFRET) enables direct observation of these dynamics in individual diffusing receptors, either freely diffusing in solution, using confocal microscopy, or immobilized on surfaces, using Total Internal Reflection Fluorescence (TIRF) camera-based microscopy. However, these modalities are limited to distinct timescales – faster than milliseconds or slower than hundreds of milliseconds, respectively – due to constraints of observation time and camera frame rate, while the slow dynamics observed in immobilized receptors may be perturbed by tethering. To overcome these limitations, we employed smFRET in combination with Anti-Brownian Electrokinetic (ABEL) trapping to extend the observation window of untethered A2A adenosine receptors (A_2A_AR) embedded in lipid nanodiscs from milliseconds to seconds. This approach enabled us to characterize the conformational heterogeneity in both apo and ligand-bound A_2A_AR, effectively bridging the gap between fast dynamics measured in freely diffusing receptors and slow dynamics observed in immobilized receptors. Our results, taken together with previous studies, demonstrate that both inactive-like and active-like conformations are populated in apo and ligand-bound A_2A_AR and dynamically interconvert. Apo and antagonist-bound receptors are predominantly locked in inactive-like or active-like states, exhibiting only slow exchange dynamics on timescales longer than hundreds of milliseconds. In contrast, agonist-binding not only shifts the receptor population toward the active-like state but also accelerates the conformational exchange by nearly three orders of magnitude: from hundreds of milliseconds to sub-millisecond timescales. These results underscore the dual role of agonists in modulating both the equilibrium and dynamics of receptor conformations. Our study highlights the power of ABEL-FRET to probe structural dynamics of GPCRs, offering valuable insights into GPCR conformational landscapes and aiding the development of more effective GPCR-targeted therapeutics.

## Introduction

G protein-coupled receptors (GPCRs) orchestrate critical physiological processes such as vision, neurotransmission, inflammation, and blood pressure regulation. Characterized by their seven-transmembrane helix topology, GPCRs constitute the largest family of membrane proteins in the human genome, encompassing over 800 members^2,3^. Their central role in human health is underscored by the fact that more than 30% of all FDA-approved drugs target GPCRs as their primary mode of action^4,5^. Many endogenous ligands and drugs bind to the extracellular ligand-binding pockets of GPCRs, triggering conformational changes in the transmembrane and intracellular regions, which alter interactions between the receptors and their intracellular signaling partners, such as G proteins and arrestins^6,7^. The structural flexibility of GPCRs underlies the wide variety of mechanisms by which ligands modulate the receptors’ activity^8–10^. For example, a GPCR can exhibit various activity levels when bound to agonists with different efficacies^11^. The activity of an agonist-bound GPCR can be further tuned by allosteric modulators that bind to additional, less evolutionary conserved pockets in the receptor’s structure^12,13^. Finally, biased agonists shift the equilibrium between various signaling pathways triggered by the same GPCR (e.g. those mediated via various types of G proteins, arrestins, or GRK kinases)^14,15^. This diversity in the pharmacological profiles of GPCR ligands opens opportunities for more effective use of GPCR-targeted drugs in clinical settings. However, the structural flexibility of GPCRs, which underlies their complex pharmacology, poses serious challenges for structure-based drug design and for the understanding of fundamental mechanisms of GPCR activation.

Despite the challenges posed by the structural flexibility of GPCRs in obtaining high-resolution structures via X-ray crystallography or cryo-electron microscopy, these techniques have significantly advanced our understanding of the structure-function relationship in GPCRs^16,17^. A notable drawback of these techniques is that they provide only static snapshots of stable conformational states, with limited information about conformational dynamics^9^. A diverse set of complementary methods, including nuclear magnetic resonance (NMR), double electron-electron resonance (DEER), mass-spectrometry, and single-molecule fluorescence microscopy, reveals structural heterogeneity and complex conformational dynamics in GPCRs^9,18^. They show that receptors switch between multiple stable states on timescales ranging from nanoseconds to seconds^9,18^. Faster motions correspond to near-equilibrium vibrations and rotations of amino acid side chains and unstructured parts of the receptor, while slower motions involve higher-order rearrangements of secondary structure elements and loops^8^. GPCR ligands, ranging from inverse to full agonists and including biased agonists and allosteric modulators, affect the rates of conformational changes in GPCRs and the relative populations of stable conformations^19^.

Single-molecule fluorescence microscopy is a powerful method for monitoring conformational dynamics in proteins and, in particular, GPCRs^20,21^. Single-molecule sensitivity of fluorescence microscopy surpasses the need for ensemble averaging, allowing the identification of distinct protein states that co-exist within an ensemble of proteins under equilibrium conditions. Conformational changes in a protein are typically monitored based on conformation-dependent fluorescence changes from a single environment-sensitive fluorescent dye (e.g. in single-molecule photoisomerization-related fluorescence enhancement (smPIFE^22^), photoinduced electron transfer (PET^23^) or a pair of donor and acceptor dyes for smFRET^24,25^. In smFRET experiments, the conformational changes of the protein affect the proximity and relative orientations of the fluorophores attached to its mobile structural elements, leading to changes in the efficiency of energy transfer from the donor to the acceptor dye (Figure 1a). FRET is extremely sensitive to variations in the mutual position of the dyes, including inter-dye distance changes at the Ångström-scale and alterations of the dye orientations, which result in measurable changes of the ratio between donor and acceptor fluorescence intensities, making this technique highly sensitive for observing conformational dynamics.

**Figure 1.**
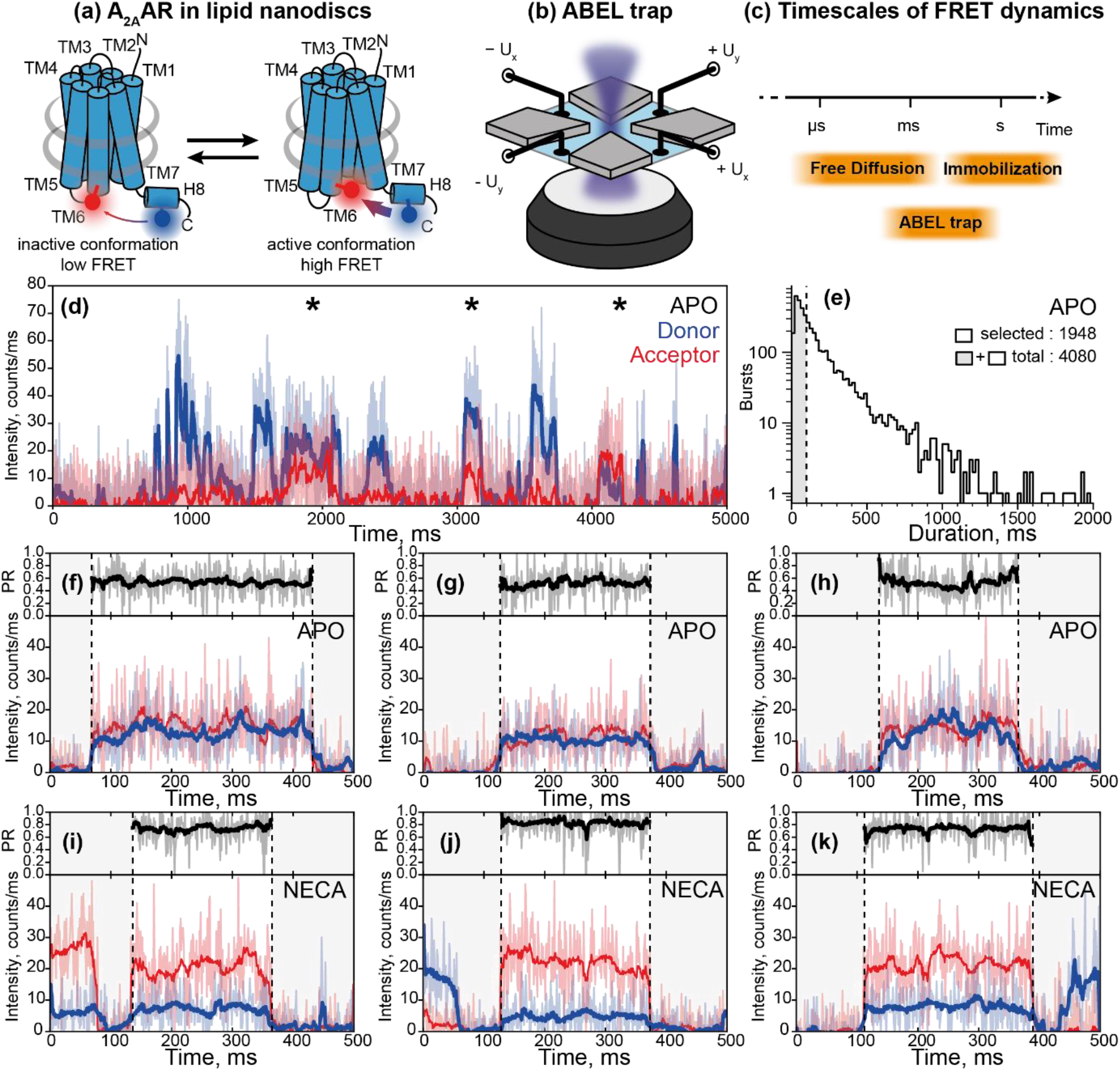
smFRET dynamics of individual A_2A_ARs in the ABEL trap. **(a)** A_2A_ARs were labeled with donor (Alexa488, blue circle) and acceptor (Atto643, red circle) fluorescent dyes at the transmembrane helix TM6 and the C-terminal helix H8 and were reconstituted in lipid nanodiscs (grey belts). The FRET efficiency in labeled A_2A_AR increases upon receptor activation. **(b)** Sketch of the ABEL trap setup with four electrodes in the PDMS-based microfluidic chamber. The FRET donor fluorophore is excited by a fast-switching laser beam pattern, and the fluorescence of both donor and acceptor on individual receptors is collected through the objective of a custom-built confocal microscope. **(c)** The timescales of the structural dynamics observed using smFRET in the ABEL trap and the most common smFRET experiment modalities, i.e., confocal microscopy with freely diffusing molecules and TIRF camera-based microscopy with immobilized molecules. **(d)** Fluorescence traces in the donor (blue) and acceptor (red) channels were recorded for individual A_2A_ARs held in solution by the ABEL trap. Background-corrected fluorescence traces were binned in 1-ms bins (thin semitransparent lines) and then smoothened using Chung-Kennedy filtering (thick lines). Fluorescence bursts with both donor and acceptor fluorescence are depicted with asterisks. **(e)** The histogram of the durations of the individual fluorescence bursts recorded for the apo A_2A_ARs. The photon bursts shorter than 100 ms were excluded from the analysis (grey area). Out of total 4080 bursts, 1948 bursts longer than 100 ms were selected and further filtered based on fluorescence intensity (see Supplementary figure 1, Supplementary table 1). **(f-k)** Exemplary fluorescence traces for the individual apo A_2A_ARs (f-h) and A_2A_ARs with agonist NECA (i-k). Donor (blue) and acceptor (red) fluorescence signals were plotted after binning with a 1-ms time-window and background subtraction (thin semitransparent blue and red lines). The areas outside of the selected fluorescence bursts are grey. For each 1-ms bin within bursts, the Proximity Ratio (PR; Equation 1) was calculated (thin semitransparent black lines), and then, PR profiles were smoothened with Chung-Kennedy filtering (thick black lines)^1^.

SmFRET experiments are typically performed in one of two modalities, depending on whether proteins freely diffuse in the sample or are immobilized on a glass surface^25^. In the first modality, freely diffusing molecules transiently cross the detection volume of a confocal fluorescence microscope. The observation time is defined by the diffusion of the molecule under investigation across the detection volume, which is typically occuring in one to a few milliseconds. In the second modality, target molecules are immobilized on a cover glass and imaged via a TIRF microscope or a scanning confocal microscope, allowing observation of slower conformational dynamics^26^. Immobilized molecules can be observed for longer periods, up to several minutes or even hours, limited mostly by photobleaching of the fluorescent dyes^27–29^. Immobilization of target molecules by tethering and their interaction with the surface, however, can potentially affect their structural dynamics, posing challenges for immobilization-based experiments^25,30,31^. Alternative approaches to observing individual fluorescently labeled molecules for longer periods without immobilization include measuring diffusion in viscous media, e.g. glycerol, or a polyacrylamide gel^32–34^, conjugating molecules to large slowly-diffusing particles or liposomes^35^, confining freely diffusing molecules within a microfluidic chip^36,37^, or tracking them, e.g. via orbital tracking^38–40^or camera-based tracking within excitation region of highly inclined and laminated optical (HiLO) excitation^32^. The diffusion of fluorescently labeled molecules in solution can also be actively counteracted using an Anti-Brownian ELectrokinetic (ABEL) trap, where untethered molecules are held in the field of view of a microscope for up to several minutes, enabling the monitoring of slow conformational dynamics all while molecules are not physically tethered^41–45^.

In an ABEL trap, molecules diffuse within a ~1-micron-thin plane of a microfluidic device and transiently enter the detection area of a scanning confocal microscope (Figure 1b). When a fluorescently labeled molecule enters this region, its position is estimated in real time based on its fluorescence emission. An external electric field - controlled by a feedback loop and applied via electrodes - is then used to counteract the molecule’s random Brownian motion and maintain it near the center of the scanned area^42^. By rapidly adjusting the applied voltages, charged molecules can be stably held within the microscope’s field of view for durations ranging from seconds to minutes—several orders of magnitude longer than in traditional diffusion-limited single-molecule experiments. During this extended observation time, emitted photons are continuously recorded, enabling detailed analysis of the molecule’s conformational dynamics. The ABEL trap has previously been used to monitor monomer-dimer transitions of GPCRs extracted from membranes using styrene–maleic acid copolymers (SMALPs), as well as conformational changes in GPCR monomers reconstituted in detergent micelles, using single-fluorophore labeling strategies^46,47^.

Here we combined ABEL trapping with smFRET to investigate the conformational dynamics of a prototypical GPCR, the human A_2A_ adenosine receptor (A_2A_AR). A_2A_AR plays an important role in the regulation of cardiovascular tonus, sleep, inflammation, and neurotransmission of dopamine and glutamate^48,49^. It is a promising target for drugs against insomnia, chronic pain, depression, Parkinson’s disease, and cancer^49–51^. A_2A_AR is one of the most studied GPCRs, with two single-particle cryo-EM (cryogenic-electron microscopy)^52,53^, two micro-crystal electron diffraction^54,55^and over 75 X-ray crystallographic structures available for the antagonist-bound^52,56,57^or agonist-bound^58,59^receptors as well as for a ternary complex of the receptor with an agonist and an engineered G protein^53,60^. NMR studies uncovered the conformational dynamics of A_2A_AR across a broad range of time scales from nanoseconds^61^to milliseconds^62–65^, and seconds^64,66^. PET-FCS^67^and smFRET^68^detected sub-millisecond conformational dynamics in freely diffusing A_2A_AR, while single-dye measurements with immobilized A_2A_AR revealed slow conformational changes over several seconds^69,70^. In this study, we observed individual A_2A_AR molecules in lipid nanodiscs for up to two seconds without immobilization using ABEL trapping and monitored their conformational dynamics and ligand-induced conformational changes using smFRET.

## Results

### ABEL-FRET measurements reveal structural heterogeneity and agonist-induced conformational changes in A_2A_AR

To observe conformational dynamics in A_2A_AR using smFRET, we fluorescently labeled and purified the recombinant human A_2A_AR (Figure 1a). For fluorescent labeling, two cysteine mutations were introduced: L225C^6.27^on the intracellular side of the transmembrane helix 6 (TM6) and Q310C^8.65^on the intracellular C-terminal helix 8 (H8, superscripts denote Ballesteros-Weinstein residue numbering in class A GPCRs^72^). These cysteines were labeled with donor and acceptor fluorophores, Alexa488-maleimide and Atto643-maleimide as in our previous study, where we used the same labeling positions and fluorescent dyes to measure sub-millisecond conformational dynamics in A_2A_AR using burst-wise smFRET with freely diffusing molecules^68^. The functional integrity of the fluorescently labeled and reconstituted in lipid nanodiscs A_2A_AR from the same purification batch was previously confirmed^73^. In the same work, we demonstrated that the FRET efficiency in the double-labeled A_2A_AR_L225C/Q310C_ increased upon receptor activation by agonists^73^.

Using an ABEL trap, we now observed individual A_2A_AR molecules typically for 0.1-1 s in the detection area of 2.34 × 2.34 μm^2^, i.e., a roughly 100-fold longer residence time (Figure 1b-k) than previously achieved with smFRET on freely diffusing A_2A_AR molecules in solution. We excited the donor fluorophore with a continuous-wave laser at 491 nm and simultaneously recorded direct emission from the donor and FRET-induced emission from the acceptor. The sharp increases in the fluorescence in both channels manifested the detection of individual double-labeled receptors, followed by sharp decreases back to the background level when the receptors escaped from the trap, or photobleaching of the donor fluorophore occurred. The receptors with absent or photobleached acceptors, where donor fluorescence was not accompanied by a FRET signal, were excluded from the analysis. If the acceptor fluorophore was bleached during the observation time, the fluorescence traces up to the moment of photobleaching were analyzed. The fluorescence intensities of individual molecules were predominantly distributed around a single peak, with no evidence of distinct subpopulations, with an average intensity of 20-30 photons/ms (sum of both color channels, averaged over the observation times; Supplementary figure 1, Supplementary table 1). To exclude a few potential oligomers or aggregates, we removed receptors with total fluorescence intensities exceeding 40 photons/ms from all further analyses.

To quantify FRET efficiency changes within the observation time of individual A_2A_AR molecules and among various molecules in the ensemble, we calculated the Proximity Ratio (PR) as follows:

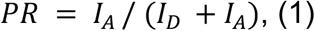

where *I*_*D*_ and *I*_*A*_ are background-corrected fluorescence intensities of donor and acceptor, averaged either within 1-ms time bins (bin-wise PR) or over the entire fluorescence burst from the individual receptor (burst-wise PR).

We compared the distributions of the burst-wise PR across different samples including A_2A_AR in its ligand-free (apo) state as well as bound to either an antagonist ZM241385 or agonists NECA (5’-N-ethylcarboxamidoadenosine) or adenosine (Figure 2b). In agreement with our previous studies with A_2A_AR labeled with the same dyes at the same labeling positions^68,73^, the antagonist did not change PR, while the agonists increased PR compared to the ligand-free condition. Under all four conditions, the PR distribution were much wider compared to what could be expected from shot-noise-limited data, indicating A_2A_AR states that do not fully interconvert during the observation times of hundreds of milliseconds (Figure 2b). These long-lived receptor’s states can be related to the slow conformational dynamics of A_2A_AR observed in the experiments with immobilized receptors^69^.

**Figure 2.**
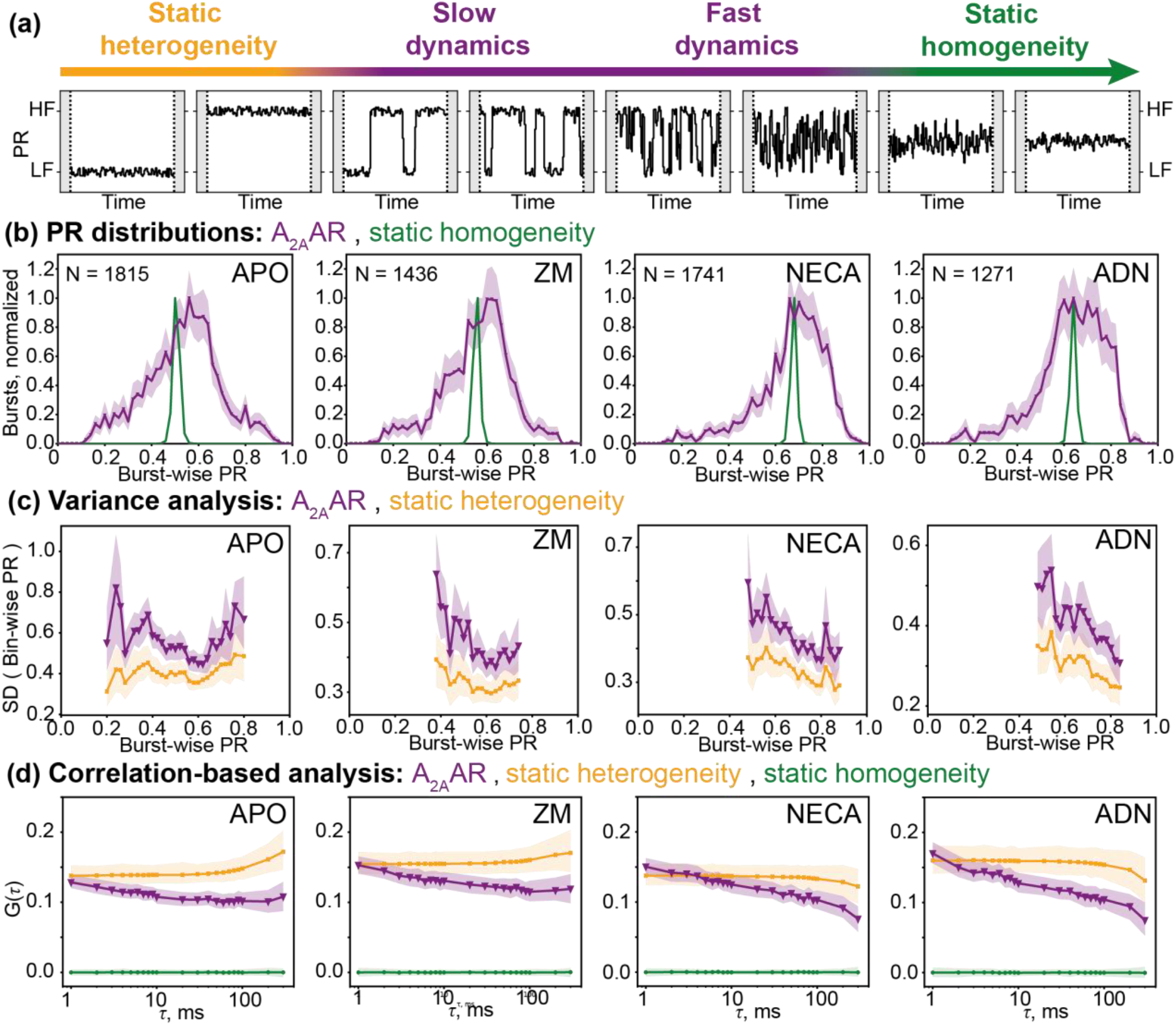
Burst-wise analysis of the structural heterogeneity and dynamics of A_2A_AR. **(a)** Schematic drawings of the PR traces expected for a molecule switching between low-FRET (LF) and high-FRET (HF) states. Left, if the switching is too slow to occur within the observation time of individual molecules, the data is indistinguishable from the “static heterogeneity” scenario, where low-FRET and high-FRET species seemingly do not interchange. Right, if the switching is fast enough to average over the binning time of the experiments, the data is indistinguishable from the “static homogeneity” scenario, where all molecules have the same PR that remains constant over time. **(b)** Experimental distributions of the burst-wise PR measured for A_2A_AR without ligands (APO, N=1948 bursts), with antagonist ZM241385 (ZM, N=1455 bursts), with agonist NECA (NECA, N=1775 bursts), and with agonist adenosine (ADN, N=1342 bursts) are shown (purple lines). Simulated distributions for the “static homogeneity” scenario are also given (green lines) to show the expected broadening of the PR distributions due to shot noise. In the “static homogeneity” simulations, the ground-truth burst-wise PR matches the ensemble-averaged burst-wise PR in the A_2A_AR data. **(c)** Standard deviations of the PRs measured in 1-ms time-bins were averaged in the groups of bursts equally spaced along the burst-wise PR axis, and were plotted for the apo and ligand-bound A_2A_ARs (purple lines) and the simulated “static heterogeneity” scenario (orange line). In the “static heterogeneity” simulations, the distribution of the PR matches those of the A_2A_AR data. PR bins with less than 20 bursts were not analyzed. **(d)** Autocorrelation functions for the PRs calculated in 1-ms time bins are plotted for the apo and ligand-bound A_2A_AR (purple lines), and for simulated “static heterogeneity” (orange) and “static homogeneity” (green) scenarios. In the simulated data, the numbers of bursts, their durations, and the levels of the shot noise match those of the experimental data for A_2A_AR shown on the same panel. The 95% confidence intervals were obtained via statistical bootstrapping (shaded colored areas)^71^.

### Variance analysis reveals moderate but significant conformational dynamics in A_2A_AR

Under all measured conditions, ligand-bound or apo, the PR fluctuations within detection times of individual receptors were significantly higher than those expected from the shot noise limit (Figure 2c). To make this comparison, we adapted the Burst Variance Analysis^74^approach to ABEL-FRET data and simulated the data for the “static heterogeneity” scenario with the same number and lengths of single-molecule fluorescence bursts, the same levels of total fluorescence signal and background, and the same PR distributions. In this simulation, each molecule was attributed an independent, constant PR, and the fluctuations were purely due to the shot noise (Figure 2a). The PR fluctuations larger than the shot noise limit indicate conformational dynamics of A_2A_AR as well as complex photophysics of the fluorophores or fluctuations of the background fluorescence.

### Correlation-based analysis reveals slow dynamics and long-lived conformational states of A_2A_AR

To analyze the time scales of the conformational dynamics observed in A_2A_AR by ABEL-FRET, we calculated the normalized autocorrelation function for the PR fluctuations as follows:

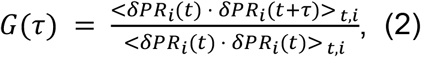

where *i* and *t* indices correspond to the individual fluorescence bursts and time bins within them, respectively, and averaging is performed over all pairs of time bins t and t+τ from all bursts. The PR fluctuations δPR_i_ (t) were calculated with a reference of PR averaged over all 1-ms time bins for all detected molecules:

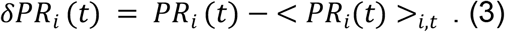

The normalized autocorrelation function equals 1 at lag time τ=0 (not shown in log-scale plots), and in essence, represents the fraction of total PR variance that remains correlated as the system’s memory of its starting state fades over time lag τ. In the apo and ligand-bound A_2A_AR, the autocorrelation is positive over the whole range of τ from the shortest binning time τ = 1 ms, to the longest τ = 300 ms (Figure 2d). The autocorrelation decreases almost linearly with log(τ) in all apo or ligand-bound conditions. The positive autocorrelation in this analysis indicates the heterogeneity within the ensemble of A_2A_ARs in different receptor states, distinguished by PR. In an imaginary scenario of “static homogeneity”, where all molecules have the same conformation and the PR fluctuations are due to the shot noise in the fluorescence data, such an analysis would results in the autocorrelation of zero for all τ>0. The zero autocorrelation would also be expected if the conformational dynamics of a protein is much faster than the 1-ms binning time used in the analysis (Figures 2a, 2d).

In our autocorrelation-based analysis, the decrease of the autocorrelation over time indicates the conformational dynamics of A_2A_AR on this time-scale. In a hypothetical scenario of a protein switching between two states with rate constants *k*_*12*_ and *k*_*21*_, the autocorrelation curve is expected to be sigmoidal with a 50%-loss of the autocorrelation at the relaxation time of τ = (k_12_ + k_21_)^−1^, and approaching zero at much longer times. Alternatively, in the case of “static heterogeneity”, where molecules have distinct static states with no inter-state switching, a positive autocorrelation that does not decrease with τ is expected (Figure 2d). Notably, the scenario of a “static heterogeneity” also describes the case, where inter-state switching occurs on much longer time-scales than the molecule’s residence times within the ABEL trap region (Figure 2a).

The low starting levels of the autocorrelation curves in our analysis can potentially be affected by the shot noise in the data, or by the fast sub-millisecond dynamics of the receptors (Figure 2d). In the apo and ligand-bound A_2A_AR, these levels are approximately 20%, indicating that 80% of the variance in the PR data is not correlated between consecutive 1-ms time bins. To test the fast conformational dynamics hypothesis, we performed computer simulations of the ‘“static heterogeneity” scenario, where the shot noise is the only source of the PR fluctuations within the individual trapping events (Figures 2a, 2d). In these simulations, we obtained similar starting levels of the autocorrelation, suggesting that the low starting values of the autocorrelation are due to shot noise rather than fast conformational dynamics.

The PR autocorrelation curves measured for the apo and ligand-bound A_2A_ARs revealed a substantial residual correlation between PRs measured with a time delay of hundreds of milliseconds, indicating long-lived receptor’s states (Figure 2d). This high residual autocorrelation further supports the “static heterogeneity” scenario, where molecules have distinct conformations that remain unchanged during trapping times, as a good approximation for A_2A_AR. The subtle but significant decrease in the PR correlation measured for A_2A_AR does not show clear sigmoidal transitions within the 1-300 ms time scale and, most probably, indicates either complex minor multi-state conformational variability on the observed time scales or the onset of slower dynamics with exchange relaxation times longer than 300 ms.

Upon structural changes in A_2A_AR, PR values vary due to alterations in the distance between the dyes or their relative orientation, resulting in true FRET changes. Additionally, FRET-independent alterations in brightnesses of individual dyes, e.g. due to changes in their local environment, can also manifest as PR variations. To disentangle the contributions of FRET changes and other processes, we analyzed correlation functions for donor and acceptor fluorescence intensities and their sum (Supplementary Fig. 2). Under all apo and ligand-bound conditions, intensities in donor and acceptor channels were anticorrelated indicating that FRET changes played a major role in the observed heterogeneity in PR values. However, the indications of FRET-independent fluorescence changes in the A_2A_AR were also evident, i.e. a decrease in the cross-correlation function G_DxA_ at 1-10 ms delay times and a decrease in autocorrelation functions without a corresponding increase in cross-correlation at 10-100 ms. These observations suggest that FRET changes differentiate the long-lived A_2A_AR states and underlie the PR variability across many molecules, while other FRET-independent processes contribute to PR fluctuations within observation times of individual A_2A_AR molecules.

### Long-lived states of A_2A_AR are confirmed by the Recurrence Analysis on the PR distributions

To further investigate the long-lived A_2A_AR states, we compared how the PR distributions change over time for the A_2A_AR molecules that at a certain time-point exhibited a PR above (“high-FRET”) or below (“low-FRET”) the ensemble-averaged median (Figures 3a, 3b). Under all four apo and ligand-bound conditions, A_2A_ARs with a higher-than-average PR in a 1-ms time bin exhibit, in general, a higher-than-average PR in the subsequent 1-ms time bin. This trend is preserved even for the time delay of 200 ms, with the PR distributions barely shifting towards the “equilibrium” of the ensemble distribution. An apo A_2A_AR molecule that showed a “high-FRET” PR in a given time bin has 65% and 59% probabilities of showing the “high-FRET” PR 1 ms and 200 ms after the initial observation, respectively (Figures 3c, 3d). Both probabilities are significantly above a random 50% value expected if the time delay between two data points is much longer than the typical timescale of the conformational dynamics.

**Figure 3.**
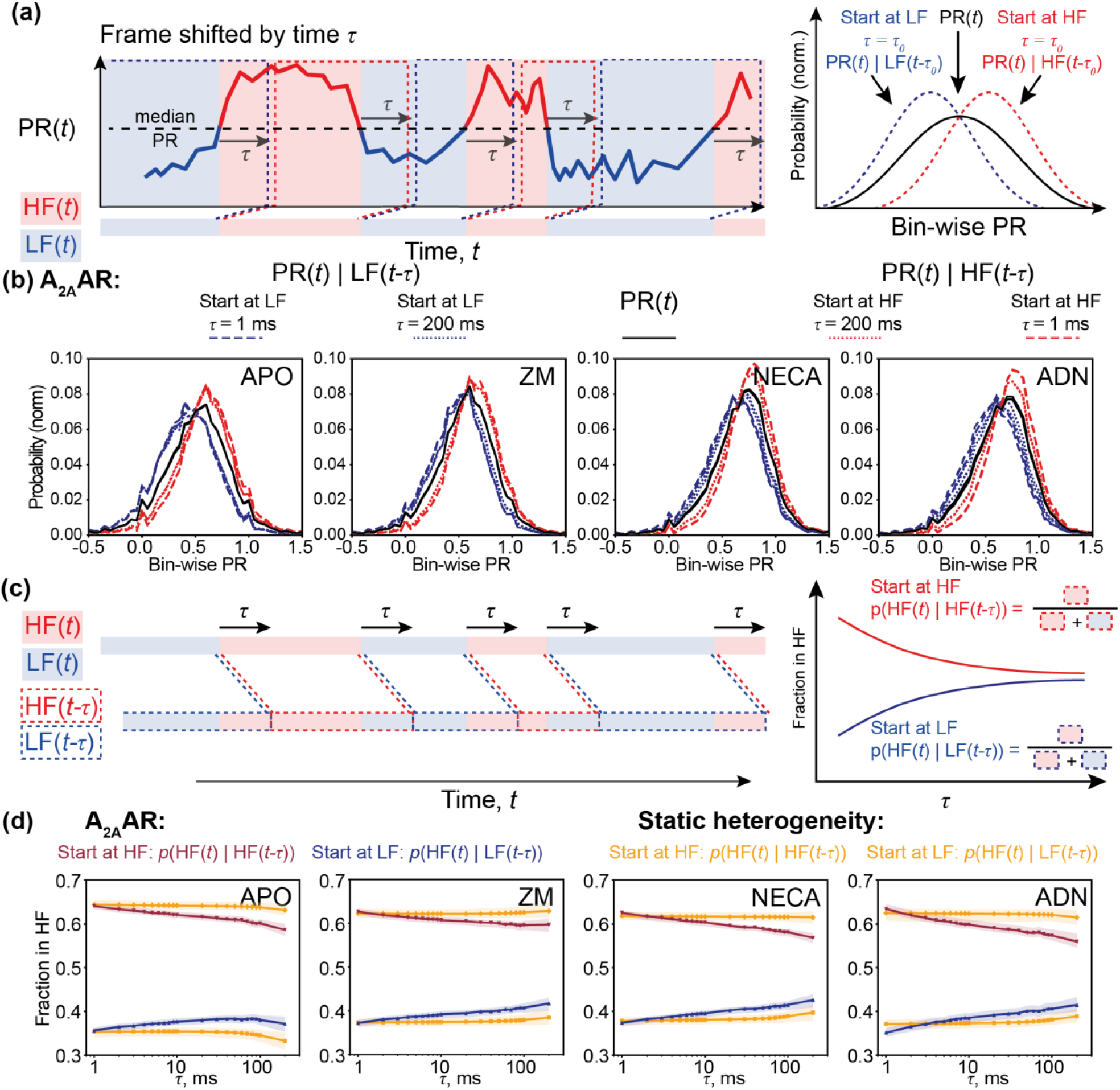
Recurrence Analysis of the PR distributions. **(a)** In the time-trace of the bin-wise PR (thick line), positive (red areas, above median PR) and negative (blue areas, below median PR) fluctuations are assigned to the high-FRET (HF) and low-FRET (LF) regions, respectively. For a given time delay *τ*, the distributions of bin-wise PR are plotted for the regions shifted by *τ* over the time axis from the HF regions (PR(*t*) | HF(*t*-*τ*), dashed red lines) or from the LF regions (PR(*t*) | LF(*t*-*τ*), dashed blue lines). The shifts of these distributions from the general bin-wise distribution of PR (black line) indicate higher than random chance of finding the protein in the same state for the time points *t*-*τ* and *t*. **(b)** Distributions of the bin-wise PR in the regions shifted from the LF (“Start at LF”) or HF (“Start at HF”) regions by time delays *τ*=1 ms and *τ*=200 ms are shown for the apo and ligand-bound A_2A_AR. The general distributions of bin-wise PR are shown in black. **(c)** The conditional probability of observing the positive fluctuation in PR at the time *t* (red areas), given that earlier, at the time *t*-*τ*, a positive fluctuation was also observed (dashed red contours), is calculated as a function *p*(HF(*t*) | HF(t-*τ*)) of *τ* (solid red line). Similarly, the probability of the positive fluctuation in PR at the time *t*, given that a negative fluctuation was observed at the time *t*-*τ* (dashed blue contours), is calculated as a function *p*(HF(*t*) | LF(t-*τ*)) of *τ* (solid blue line). On the right, two curves converge to the same value at times much longer than the inter-state exchange relaxation time of the target molecules. **(d)** Two conditional probabilities of the positive fluctuation in PR are plotted for the apo and ligand-bound A_2A_AR. Two orange lines show the same curves for the simulated “static heterogeneity” scenario (*p*(HF(*t*) | HF(t-*τ*)) - higher line; *p*(HF(*t*) | LF(t-*τ*)) - lower line). The 95% confidence intervals were obtained via statistical bootstrapping (shaded colored areas)^71^.

The probability of observing “high-FRET” PR after it was once observed significantly decreases with the time delay between two data points for the apo and ligand-bound A_2A_AR (Figure 3d). This decrease indicates moderate but significant conformational dynamics of A_2A_AR on the 1 to 200 ms time range. For reference, in simulations of the “static heterogeneity” scenario, where molecules have distinct conformations that remain unchanged over the trapping time, and the PR variations are due to the shot noise in the fluorescence data, the probability of recording a “high-FRET” PR 1-ms after it was once recorded is similar to what we observe for A_2A_AR, but this probability does not decrease for the longer time delays *τ*. Notably, the similar starting levels in the experimental data and simulations indicate that the shot noise in the data renders the sub-millisecond fast receptors’ dynamics invisible in our experimental ABEL trap setup.

In summary, our findings demonstrate that A_2A_AR exhibits only moderate deviation from the “static heterogeneity” scenario, with a high probability of detecting the receptor in the same “state” in the beginning and the end of the trapping time of hundreds of milliseconds. Using the ABEL-FRET, we bridged the gap between previous studies using confocal smFRET with freely diffusing receptors^67,68^, where the sub-millisecond conformational dynamics were investigated, and the TIRF-based measurements with the immobilized receptors^69,70^, shedding light on the dynamics on the time scale of seconds. In this study, the fast dynamics of A_2A_AR could not be resolved due to the shot noise in the fluorescence data and the microsecond motion of the laser focus in the ABEL trap, and were averaged out within 1-ms time binning. The slow dynamics contributed to subtle yet statistically significant deviations from the “static heterogeneity” scenario observed in our data. We did not observe any characteristic dynamics in A_2A_AR on the time scales from milliseconds to hundreds of milliseconds - neither the autocorrelation function nor the probability of detecting the same-sign PR fluctuation showed a sigmoidal decrease within the observed time scale.

## Discussion

Using the ABEL-FRET approach, we monitored the conformational changes in the individual A_2A_ARs reconstituted in lipid nanodiscs during trapping times of 0.1 to 2 s (Figure 1). In previous ABEL trap studies, two GPCRs, the β_2_-adrenergic receptor (β_2_AR^46^) and the neurotensin receptor 1 (NTSR1^47^), were reconstituted in detergent micelles or in SMALPs, respectively, and trapped in solution for up to a few seconds. In addition to GPCRs, another membrane protein, F_o_F_1_-ATP synthase, was previously FRET-labeled and reconstituted in proteoliposomes for ABEL-FRET^75–77^ yielding comparable observation times of a few seconds for the conformational dynamics in the 10 to 100 ms time range. The nanosecond photophysics and energy transfer of other membrane proteins studied in the ABEL trap include the light harvesting complex LH2 in detergent micelles^78^. Lipid nanodiscs with a controlled diameter and lipid composition provide a native-like lipid bilayer environment and maintain functionality and high stability for a variety of reconstituted protein targets^79^. Lipid nanodiscs have been widely used for studying membrane protein structure and function^80^(e.g. via NMR, double electron-electron resonance (DEER) spectroscopy, or cryo-EM) and for drug screening^81^, therefore their temporary immobilization by ABEL trapping opens the way for many cross-method correlational studies.

Fast sub-millisecond conformational dynamics of A_2A_AR were previously observed by smFRET^68^, PET-FCS^67^, and NMR^62,63,65^, with A_2A_AR reconstituted in lipid nanodiscs or detergent micelles. In this study, we could not observe the conformational dynamics of the A_2A_ARs on this time-scale, because within 1 ms only few fluorescence photons were detected from individual receptors, i.e., about 20 to 30 photons (Supplementary Figure 1, Supplementary table 1). Thus, to counteract the shot noise, we used the 1-ms binning in the smFRET analysis, and the sub-millisecond dynamics were averaged within the individual time bins.

The long-lived structural states of A_2A_AR with dwell times longer than milliseconds were observed in previous NMR^63–66,82–84^and smFRET experiments^67,68^. In the smFRET-based studies^67,68^, the individual A_2A_ARs were observed for several milliseconds until the freely-diffusing receptors left the focal spot of the microscope. With the ABEL trap, we extended the observation time of the individual receptors to 2 s (Figure 1e). On this timescale, we see only minor changes of the FRET efficiencies in the individual A_2A_AR molecules, with a high probability of the molecule to remain in the same state throughout the photon burst (Figures 1f-k, Figures 2c, 2d, 3c, 3d). Thus, we can update the previous estimates of the dwell times for the long-lived A_2A_AR states observed by smFRET by two orders of magnitude, from milliseconds to hundreds of milliseconds. The existence of long-lived states with dwell times exceeding 100 ms is supported by the recent NMR-based study with A_2A_AR reconstituted in lipid nanodiscs^84^. The minor, yet significant, dynamics on the 1 to 300 ms time scales that we observed using the correlation-based analysis (Figure 2d) and the recurrence analysis (Figures 3c, 3d) of the PR distributions can be related to the sub-second dynamics of the A_2A_ARs previously measured via NMR^64,66^ or is an onset of even slower dynamics.

Using molecular dynamics simulations, we previously showed that the FRET changes observed upon the activation of the A_2A_AR_L225/Q310C_ can be due to the outward movement of the intracellular tip of TM6, followed by the movement of the dye attached to L225C^6.27^toward the G protein binding cavity^68^. Using the ABEL trap, we observed long-lived FRET-states of A_2A_AR with dwell times exceeding 200 to 300 ms (Figure 2d, Figure 3d). We hypothesize that these states represent slow structural rearrangements of the intracellular part of the receptor around the major active and inactive conformations. These slow rearrangements might be functionally important for the recognition of the G protein and can be linked to the slow minutes-long rearrangements in the structure of G protein bound to A_2A_AR, which was previously observed via HDX MS^85^.

The slow seconds-long switching between the A_2A_AR conformations was previously monitored using single-molecule fluorescence microscopy with up to 0.1 s temporal resolution using immobilized A_2A_ARs reconstituted in nanodiscs^69,70^. The long-lived states observed in our experiments can be related to these previously reported states. The direct comparison between the two single-molecule studies is challenging because different strategies were used to follow the conformational dynamics of the receptor. In the studies of Wei et al.^69,70^, the structural changes around the intracellular tip of TM7 were probed with the environment-sensitive dye Cy3, while in our study the movement of TM6 was probed via the FRET pair of dyes on TM6 and the C-terminal helix H8 (Figure 1a). We hypothesize that the same global conformational rearrangements in A_2A_AR, occurring on the time-scales of seconds and involving both TM6 and TM7, influence the fluorescence readouts in our study and in the study of Wei et al^69,70^.

Using new insights from the ABEL-FRET, we update the action model of the A_2A_AR that we introduced in our previous smFRET study^68^. As in the initial three-state model, the active-like state with the highest FRET efficiency (HF) is the most populated in agonist-bound A_2A_AR and the inactive-like state with the intermediate FRET efficiency (MF) is the most populated in apo and antagonist-bound A_2A_AR (Figure 4). The previously observed A_2A_AR state with the lowest FRET efficiency (LF) was the least populated under all apo or ligand-bound conditions and was assigned to non-functional or improperly folded receptors. In our ABEL-FRET measurements, we directly excited only donor, but not acceptor fluorophores. Therefore the molecules with low FRET efficiencies were indistinguishable from those without active acceptor dyes and were not analyzed. Since the LF state was the least populated state, which likely does not describe a functional receptor, and was not observed in ABEL-FRET experiments, we did not include it in our updated model. We showed that agonist-bound A_2A_AR switches between MF and HF states with the exchange time of ~0.4 ms, while in apo and antagonist-bound A_2A_AR this switching is slower (>100 ms). We use a schematic two-dimensional energy landscape to illustrate this fast agonist-induced switching between MF and HF together with long-lived heterogeneities within individual states revealed via ABEL-FRET. In this updated A_2A_AR energy landscape, the inter-state energy barrier between MF and HF is low in agonist-bound A_2A_AR and high in the apo and antagonist-bound A_2A_AR, and the intra-state energy barriers among the MF and HF sub-states are high under all apo and ligand-bound conditions.

**Figure 4.**
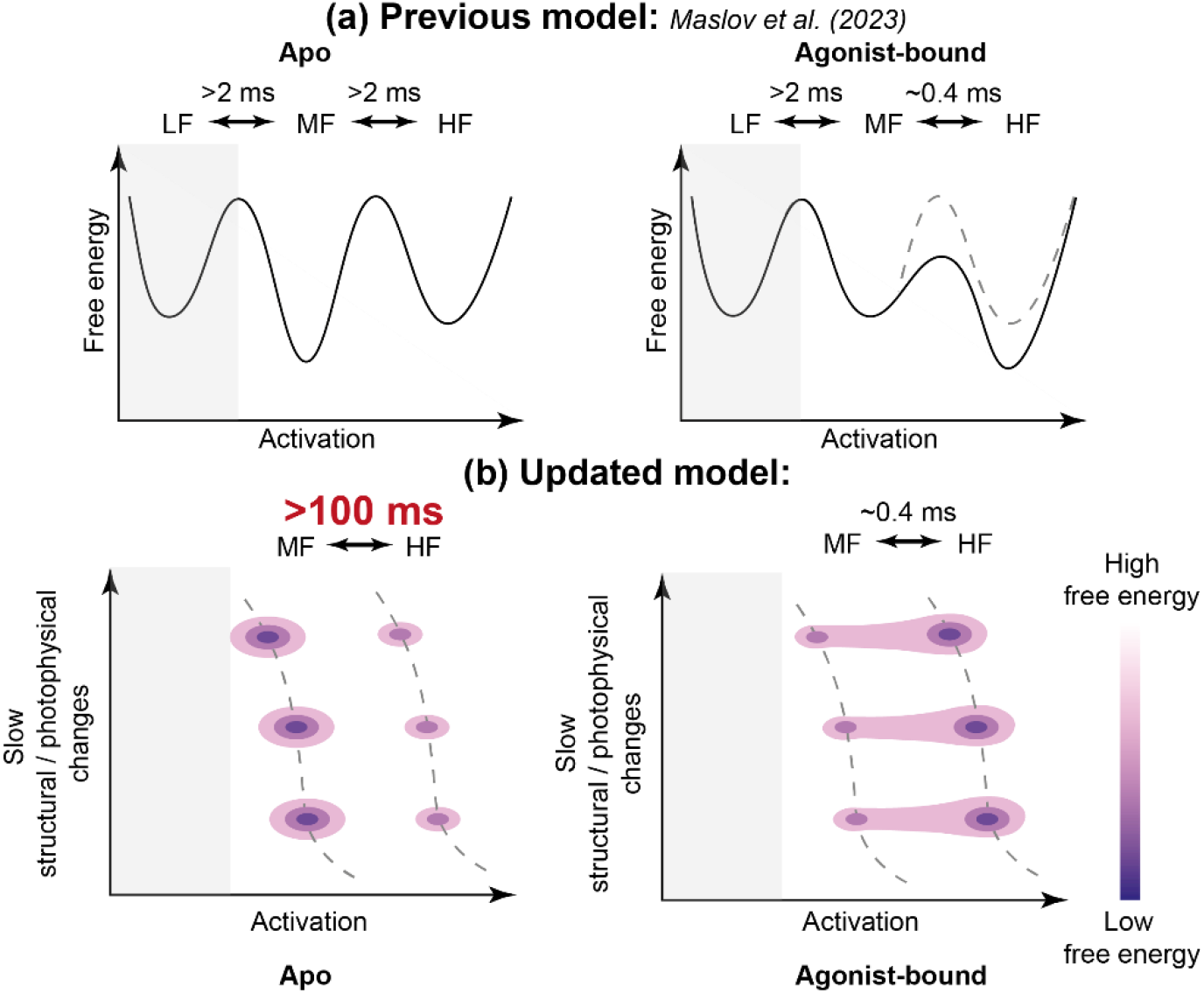
Updated action model of the A_2A_AR. **(a)** The three-state action model of the A_2A_AR proposed in ref. ^68^, and corresponding energy landscapes for the apo and agonist-bound A_2A_AR. **(b)** Updated scheme of two-dimensional energy landscapes shows long-lived sub-states within inactive-like state with intermediate FRET efficiency (MF) and active-like state with high FRET efficiency (HF). We did not include a state with the lowest FRET efficiency (LF, grey areas) in our updated model as it was the least populated and assigned to non-functional or improperly folded receptors in the previous study^68^and, in the current study, it was not observed as it was indistinguishable from molecules without active acceptor dyes. We updated the estimate of exchange time between MF and HF in apo A_2A_AR from >2 ms proposed in ref. ^68^ to >100 ms.

Our study showed that the ABEL-FRET approach can be used for the studies of the conformational dynamics of membrane proteins reconstituted in lipid nanodiscs. ABEL-FRET measurements bridge the gap in the time scales, where conformational dynamics in the A_2A_AR were previously monitored by single-molecule fluorescence microscopy, i.e., from 1 ms, the near-maximum time observed in the measurements with freely diffusing molecules^67,68^, to 100 ms, the shortest temporal resolution of the previous study with immobilized A_2A_AR^69,70^. Our measurements revealed no pronounced conformational dynamics in this 1 to 300 ms region thus refining the estimate for the dwell times of the long-lived states previously observed in the freely-diffusing receptors, and linking them with states observed in immobilized receptors with dwell times of seconds.

Several technical advancements hold promise for enhancing both the temporal resolution and the maximum trapping duration in future ABEL-FRET studies, thereby broadening the accessible time-scale for monitoring the structural dynamics of GPCRs and other membrane proteins. For example, the use of all-quartz ABEL trap microfluidic devices can significantly reduce background fluorescence^86^, while self-healing fluorescent dyes^87,88^or DyeCycling approaches^29^improve dye photostability and effective brightness, thus extending observation times. Furthermore, multiplexed readouts - such as fluorescence lifetime and anisotropy measurements, or assessments of particle diffusion coefficients and electrokinetic mobility - can uncover receptor states that remain indistinguishable using conventional FRET metrics^43,89,90^. Collectively, these innovations highlight the strong potential of ABEL-FRET for comprehensive, high-resolution studies of membrane protein conformational dynamics.

## Methods

### A_2A_AR sample preparation

The double mutant A_2A_AR_L225C/Q310C_ sample used here was prepared and functionally characterized in our previous study^56^. In that study, the human ADORA2A gene encoding A_2A_AR (2-316 aa) with N-terminal hemagglutinin signal peptide (MKTIIALSYIFCLVFA), FLAG-tag (DYKDDDDK), and a linker (AMGQPVGAP), mutations L225C^6.27^and Q310C^8.65^, and C-terminal 10 X His-tag was expressed in *Spodoptera frugiperda* Sf9 cells. The protein was labeled with Atto643-maleimide and Alexa488-maleimide in crude membranes, solubilized in DDM/CHS (n-dodecyl-b-D-maltoside/cholesterylhe misuccinate), purified on the TALON resin, and reconstituted in lipid nanodiscs (MSP1D1, POPC:POPG 7:3). The sample in 25 mM HEPES pH 7.5, 150 mM NaCl was supplemented with 40% glycerol, aliquoted, flash-frozen in liquid nitrogen, and stored at −80°C. The purity and homogeneity of the sample were confirmed by SDS-PAGE and analytical SEC. The labeling efficiencies were 16 % for Alexa-488 and 12 % for Atto 643. The ligand-induced changes in the thermal stability of the receptor and FRET efficiency were demonstrated via thermal shift assay (TSA) and smFRET with freely diffusing molecules as shown in ref. ^73^.

### Custom-designed confocal ABEL-FRET setup

All smFRET ABEL trap experiments were conducted on a modular home-built confocal microscope system as described before^41,47,75–77^. A PDMS-glass sample chamber was held on a piezo-driven stage (P-527.3CD, Physik Instrumente, Germany) incorporated into an inverted microscope (IX71, Olympus). A continuous-wave 491 nm laser (Calypso, Cobolt, Sweden) was set to a 32-point Knight’s tour laser focus pattern at a 8 kHz scanning speed per full pattern by two electro-optical beam deflectors (EOD, M310A, Conoptics, USA). The laser power was attenuated to 34 µW in the focal plane. The size of the laser focus pattern in the focal plane was 2.34 × 2.34 μm^2^.

The excitation and detection beam path both passed through an oil immersion objective (60x, PlanApo N, oil, NA 1.42, Olympus, Germany). A dichroic mirror (zt 488 RDC, AHF, Germany) separated the detected fluorescence from the excitation laser. A pinhole (200 µm) in the image plane of the tube lens optimized the signal-to-noise ratio. A dichroic beam splitter (HC BS 580i, AHF) spectrally separated the donor and acceptor fluorescence signals before two single photon counting avalanche photodiodes (APD, SPCM-AQRH 14, Excelitas, USA). A bandpass (ET 535/70, AHF, Germany) and a longpass (LP 594, AHF, Germany) filter further determined the precise wavelength detection range for the donor and acceptor channels, respectively. A time-correlated single-photon counting (TCSPC) module (SPC-180NX with router HRT-82, Becker&Hickl, Germany) recorded arriving photons in the FIFO mode.

A field programmable gate array module (FPGA, PCIe-7852R, 80 MHz operation frequency, National Instruments, USA) operated high voltage controllers (7602M, Krohn-Hite Corporation, USA) of the EODs and a feedback voltage amplifier (built in-house, |VPP| = 20 V) to deliver electric fields via four platinum electrodes counter-acting the molecule’s Brownian motion. An implemented Kalman filter correlated the detected photon signals to the scanning pattern positions and estimated the particle trajectory. The original control software (Labview, National Instruments) was provided by A. Fields and A. E. Cohen^41^and modified^47,76^to match our hardware configuration and to implement the Knight’s tour pattern.

### ABEL-FRET measurements

Disposable microfluidic chambers for the ABEL trap were prepared from structured PDMS and glass as described in ref. ^75^ using the published design on a borosilicate glass wafer^76,91^. PDMS with a ratio of 1:10 polymer:hardener was cured for 48 h at 50°C (Sylgard 184 elastomer, Dow, Midland, MI, USA). Holes for the electrodes in the PDMS part were punched using a sharpened needle. PDMS chips were sonicated in pure acetone, rinsed with deionized water and dried in a nitrogen stream. PDMS and 32 × 24 mm cover glasses with defined thickness (#1.5) were plasma-etched for 90 seconds and then bonded for 3 min to form a sample chamber.

Aliquots of labeled A_2A_AR_L225C/Q310C_ reconstituted in nanodiscs were thawed on ice, and the receptor was diluted in a buffer containing 10 mM NaCl, 20 mM HEPES pH 7.5 to a final concentration around 50 pM. Ionic strength of the buffer was kept low to avoid screening of electrical charge and ensure effective performance of ABEL trap. Reduced salt concentration (10 mM NaCl) was chosen to minimize electrostatic screening and thus to improve ABEL trap performance. For ligand binding, receptors were pre-incubated on ice with 10 µM ZM24138, 10 µM NECA, or 100 µM adenosine for 10 min. We filled 10 μL of the analyte solutions into the PDMS/glass chamber of the ABEL trap. The sample chamber was exchanged with fresh sample every 25 min. The measurements were performed for 50 or 100 min, respectively, for each apo or ligand-bound condition.

### Burst selection

We analyzed the time traces of the fluorescence intensity in the donor and acceptor channels using a customized version of the “Burst Analyzer” software^92^ (Becker & Hickl) with a 1-ms time binning. Fluorescence bursts were manually marked using stepwise rises of 25 to 40 counts per millisecond in the total fluorescence count rates at the beginning of the successful trapping events and stepwise drops back to the background fluorescence level at the ends of the trapping events. The mean background levels were around 10 to 15 counts per millisecond for the donor channel and 20 to 25 counts per millisecond for the acceptor channel caused by luminescence from PDMS and the cover glass.

To exclude the receptors without acceptor dyes or with photophysically inactive acceptors, only the photon bursts showing step-wise rise and decrease in the acceptor channel above the background level were selected. The acceptor blinking and bleaching events within a burst manifested in sharp decreases of the fluorescence in the acceptor channel to the background level - in these cases, the bursts were split in parts with the uninterrupted acceptor fluorescence. The background had to be subtracted for each photon burst individually because the background decreased on both channels in a time-dependent manner.

After the manual burst selection, bursts were removed from the subsequent analysis if the fluorescence intensity, combined from the two detection channels and averaged over the burst duration, exceeded 40 photons/ms or if a burst was shorter than 100 ms.

### Computer simulations and statistical bootstrapping

We used *in silico* photon re-coloring to simulate burst data under two scenarios: “static homogeneity” and “static heterogeneity”. In both cases, PR remains constant over time within each burst, and the numbers of bursts, their durations, and the shot noise levels mirror those observed in the experimental data. In the “static homogeneity” scenario, all bursts share the same burst-wise PR that matches the average PR across all 1-ms time-bins in the experimental data. In the “static heterogeneity” scenario, the distribution of the burst-wise PR matches those of the experimental data.

For each experimental burst, we simulated its “static homogeneity” and “static heterogeneity” re-colored versions that have the same number of 1-ms bins. In each k-th bin of a simulated burst, the total number of photons I(k) equals the total uncorrected photon count in the corresponding k-th bin of the experimental burst. The assignment of photons to donor or acceptor channels is then randomized according to the scenario. We re-colored the background photon counts, assuming a constant total number of background photons across all 1-ms bins within a burst. For background photon counts, the number of photons in the donor and acceptor channels were simulated as a binomially distributed variable, based on the ratio of background fluorescence intensities in the acceptor and donor channels in a given burst, determined during burst marking. For background-corrected photon counts, the number of photons in the donor and acceptor channel was simulated as a binomially distributed variable according to the ground-truth PR of the simulated burst, which was determined by the chosen scenario. In the “static homogeneity” scenario, the ground-truth PR for each simulated burst equals the mean PR averaged across all 1-ms bins in all experimental bursts. In the “static heterogeneity” scenario, it equals the PR measured in the burst before re-coloring. After re-coloring, the fluorescence intensities of the simulated bursts were corrected for the constant background and analyzed alongside the experimental bursts.

We estimated the 95% confidence intervals in the burst data analyses using statistical bootstrapping. Each experimental set of N bursts was resampled 200 times, with each resampling involving random drawing with replacement, allowing some bursts to be selected multiple times, while others might not be selected at all. For each condition (apo and ligand-bound), we analyzed the re-sampled datasets individually and determined the mean values and 95% confidence intervals for the values determined in each analysis approach.

### The variance analysis of the ABEL-FRET data

Our variance analysis builds on the approach of Burst Variance Analysis^74^, which we adapted for the analysis of ABEL-FRET data. For each burst, the standard deviation of the bin-wise PRs (*σ*) was calculated. Bursts were grouped by burst-wise PR in N = 51 equally spaced intervals (PR_i_, PR_i+1_) with centers at PR=0.00, 0.01, 0.02, … 0.99, 1.00. Only groups with > 20 bursts were analyzed. Within each group, the mean value of *σ (<σ>)* was determined and plotted against the PR values in the centers of the intervals 0.5^*^(PR_i_+ PR_i+1_). The analysis was performed for the experimental data and the simulated data for the “static heterogeneity” scenario. Statistical bootstrapping was performed to estimate the 95% confidence intervals of <*σ>*.

### The correlation-based analysis of the ABEL-FRET data

We calculated the correlation functions *G*^AxA^(*τ*), *G*^DxD^(*τ*), *G*^DxA^(*τ*), and *G*^sum x sum^(*τ*), for the fluorescence intensities of donor (D), acceptor (A), or their sum, all measured in 1-ms time-bins:

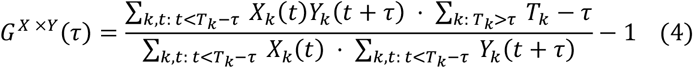

The burst width is denoted as *T*_*k*_, the X_k_(t) and Y_k_(t + τ) are values of correlated variables X and Y in the k-th burst at times t and t+*τ* from the start of that burst. Statistical bootstrapping was performed to estimate the 95% confidence intervals of G^X×Y^(τ).

We also calculated the autocorrelation function g^PR×PR^(τ) for the PR values measured in 1-ms time-bins. To compare the correlation curves between the experimental data and the simulated data for the “static heterogeneity” and “static homogeneity” scenarios, we normalized the autocorrelation functions with the standard deviations of PR:

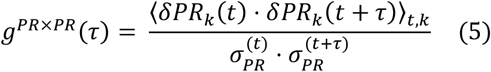

δPR_k_ (t) and δPR_k_ (t + τ) are the deviations of the PR in the k-th burst at times t and t+*τ* from the start of that burst from their ensemble-averaged mean values:

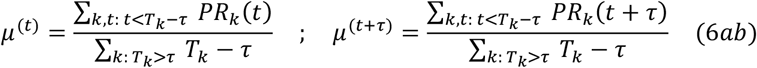

The standard deviations are calculated as:

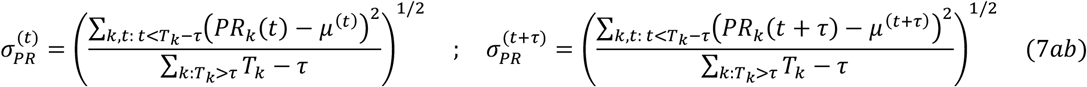

The numerator in Eq. 5 is calculated as:

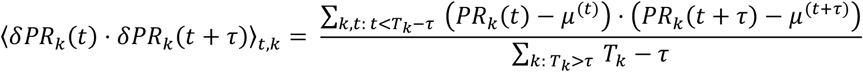

As a Pearson correlation coefficient for PR_k_(t) and PR_k_(t+*τ*), the normalized autocorrelation function g^PR×PR^(τ) can have values from −1 to 1, and equals zero if PR(t) and PR(t+*τ*) are not correlated. g^PR×PR^(τ) functions were calculated for the experimental data and for the simulated data for the “static heterogeneity” and “static homogeneity” scenarios. Statistical bootstrapping was performed to estimate the 95% confidence intervals of g^PR×PR^(τ).

### The Recurrence Analysis of PR distributions

Our Recurrence Analysis builds on the approach of Recurrence Analysis of Single Particles^93^, which we adapted for the analysis of the ABEL-FRET data. For each burst dataset, PR values were calculated in 1-ms time-bins. Here, PR_*i*_(*t*) represents the PR value in the i-th burst at time t from the start of that burst. 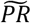 denotes the median value of the PR_*i*_(*t*) across all time-bins in all bursts. The time-bins, where 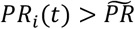 were named “high FRET” regions (HF), and time-bins, where 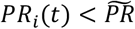 were named “low FRET” regions (LF).

For the time-lags *τ*=1 ms and *τ*=200 ms, we histogrammed the values PR_i_(t) across all bursts longer than *τ* (i: ω_i_ > τ) and all time-points t, where both t-τ and t are within the burst edges (t: 0 ≤ t − τ, t ≤ ω_i_) and PR_*i*_(*t-τ*) belongs either to LF-region (Start at LF: 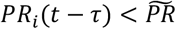) or to HF-region (Start at HF: 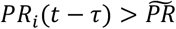 For reference, we also histogrammed the PR_i_(t) values across all bursts and all time-points.

Further, for various time-lags *τ*, we calculated the conditional probabilities of observing PR_i_(t) in the HF region 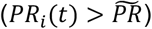, given that PR_i_(t−τ) belonged to the either LF region (Start at LF: 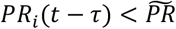) or to HF region (Start at HF: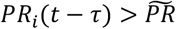). These two probabilities were plotted as functions of τ.

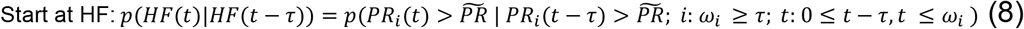

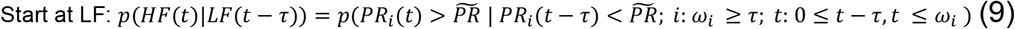

The analysis was performed for the experimental data and for the simulated data in the “Static heterogeneity” scenario. Statistical bootstrapping was performed to estimate the 95% confidence intervals of p(HF(t)|HF(t − τ)) and p(HF(t)|LF(t − τ)).

## Data availability

All data supporting this study’s findings are available upon request from the corresponding author(s). The burst data from ABEL-FRET experiments are available in Zenodo with the identifier DOI: 10.5281/zenodo.16091080.

## Code availability

Marking of the fluorescence bursts was performed in the “Burst Analyzer” software (Becker & Hickl). Further analysis was performed via custom Python scripts available in Zenodo with the identifier DOI: 10.5281/zenodo.16091080.

## Author contributions

M.B. prepared the instrumentation for the ABEL-FRET measurements

I.M., M.B., and T.G. collected ABEL-FRET data

I.M. and T.G. performed burst marking under the supervision of M.B.

I.M. analyzed the ABEL-FRET data and produced the first draft of the manuscript under the supervision of J.Ho., J.He., V.B., M.B., and T.G.

I.M., V.B., M.B., and T.G. conceptualized the study.

All the authors contributed to analyzing data, writing the original draft, reviewing, and editing.

## Acknowledgments

The authors acknowledge Dr. Polina Khorn (Moscow Institute of Physics and Technology, MIPT), who performed protein expression, labeling, purification, and nanodisc reconstitution under supervision of Dr. Aleksey Mishin and V.C. I.M. acknowledges BOF UHasselt (BOF21BL11) and funding from the European Union under the HORIZON TMA MSCA Postdoctoral Fellowships action (project MemProDx, 101149735). J.He acknowledges the Research Foundation Flanders (FWO, grant number G0B9922N). M.B. gratefully acknowledges ABEL trap funding by the Deutsche Forschungsgemeinschaft (DFG) through grants BO1891/10-2, BO1891/15-1, BO1891/16-1, BO1891/18-2. Additional support for the ABEL trap was provided by an ACP Explore project within the ProExcellence initiative ACP2020 from the State of Thuringia to the Abbe Center of Photonics (Jena).

