## Supplementary information for "ABEL-FRET bridges the timescale gap in single-molecule measurements of the structural dynamics in the A_2A_ adenosine receptor"

Maslov et al.

### Contents:

**Supplementary figure 1.** Analysis of fluorescence intensities of individual apo and ligand-bound $A_{2A}$AR molecules.
**Supplementary table 1.** Fitting parameters of fluorescence intensity distributions for apo and ligand-bound $A_{2A}$AR.
**Supplementary figure 2.** The correlation functions for the fluorescence intensities of donor and acceptor fluorophores in apo and ligand-bound $A_{2A}$AR.

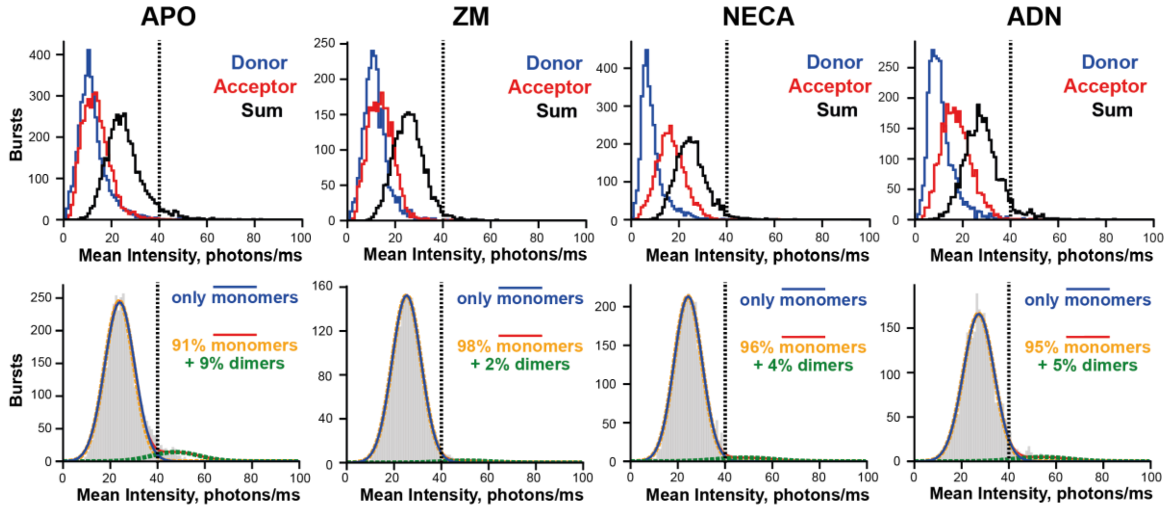


**Supplementary figure 1. Analysis of fluorescence intensities of individual apo and ligand-bound $A_{2A}$AR molecules. (A)** Histograms for fluorescence intensities in two channels and their sum, all averaged over the observation times of individual ABEL-trapped receptors. **(B)** One-Gaussian (blue lines, “only monomers”) and two-Gaussian (red lines) fits of the histograms for the sum intensity in two channels averaged over the observation times (grey stairs). Individual components of two-Gaussian fits have 2:1 intensity ratio; higher and lower intensities correspond to “dimer” (green lines) and “monomer” (orange lines) fractions, respectively. The fitting parameters are given in the Supplementary table 1. The threshold of 40 photons/ms (dashed black line) was used to exclude potential oligomers and aggregates in the subsequent analyses.

| | Monomers only | Monomers + dimers | | | |
|---|---|---|---|---|---|
| | I monomers, photons/ms | F monomers, % | F dimers, % | I monomers, photons/ms | I dimers, photons/ms |
| APO | 24±6 | 91 | 9 | 24±6 | 48±10 |
| ZM | 25±6 | 98 | 2 | 25±6 | 51±10 |
| NECA | 24±6 | 96 | 4 | 24±6 | 49±12 |
| ADN | 28±7 | 95 | 5 | 27±7 | 54±11 |

**Supplementary table 1. Fitting parameters of fluorescence intensity distributions for apo and ligand-bound $A_{2A}$AR.** Fluorescence intensities in individual $A_{2A}$AR molecules were summed over donor and acceptor channels and averaged over the molecule's observation times. In one-Gaussian ("Monomer only") and two-Gaussian ("Monomers + dimers") fits, the intensity values (*I,* photons/ms) for individual components are given as mean ± SD. In the two-Gaussian fit, individual components have 2:1 intensity ratio; higher and lower intensities correspond to "dimer" (green lines) and "monomer" (orange lines) fractions, respectively. The fractions of molecules in each component (*F*, %) are also given.

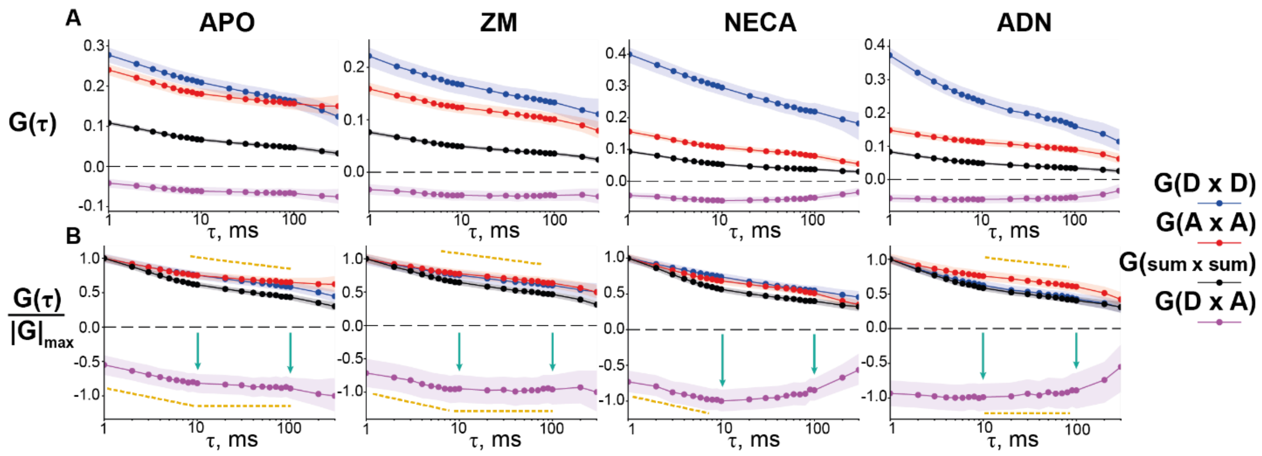


**Supplementary figure 2. The correlation functions for the fluorescence intensities of donor and acceptor fluorophores in apo and ligand-bound $A_{2A}$AR.** Autocorrelation function for donor ($G_{DxD}$) and acceptor ($G_{AxA}$) intensities, their cross-correlation function ($G_{DxA}$), and autocorrelation function for the total intensity in two color channels ($G_{sumxsum}$) are plotted against time lag $\tau$. The curves are shown without normalization (A) and after normalization to the largest absolute deviation from zero (B). The 95% confidence intervals were obtained via statistical bootstrapping (shaded colored areas)[71]. Overall anticorrelation between donor and acceptor intensities and FRET-independent trends in the data are highlighted with green arrows and yellow lines, respectively (see main text for more details).
